# Feeling the noise: Ascidians detect substrate-borne vibrations, not pressure waves

**DOI:** 10.1101/2025.08.19.671039

**Authors:** Til Böttner, Lukas Hessel, René Ortmann, Wolfgang H. Kirchner, Stefan Herlitze, Mareike Huhn

**Affiliations:** General Zoology and Neurobiology, Ruhr University Bochum, Bochum, Germany; European Molecular Biology Laboratory, Hamburg Unit, Hamburg, Germany; Behavioural Biology and Biology Education, Ruhr University Bochum, Bochum, Germany

## Abstract

Marine anthropogenic noise pollution has risen in recent decades, driving interest in its ecological effects. While research has focused on mammals and fish, marine invertebrates remain largely understudied despite their ecological importance. This study explored sound perception in the benthic ascidian *Halocynthia papillosa*, assessing whether it detects acoustic signals via waterborne pressure waves or substrate vibrations. Field and laboratory experiments exposed *H. papillosa* to frequencies from 50–1500 Hz. Contraction responses occurred at 100 Hz, 200 Hz, and 600 Hz when played through a waterproof speaker that produced both, sound pressure and substrate vibration. In a controlled laboratory setup, only substrate vibrations triggered contractions, while pure sound pressure up to 130 dB re 1 µPa elicited no response. Repeated exposure led to habituation, reducing contraction amplitude and relaxation time. These findings indicate that ascidians rely on mechanoreception, underscoring the overlooked impact of substrate vibrations from marine noise and the necessity to reconsider study design when investigating effects of marine noise on benthic organisms.

## 1. Introduction

In recent decades, anthropogenic underwater noise pollution has raised increasing concerns for marine life (Chahouri, Elouahmani, and Ouchene 2022; NRC 2003) with ambient noise levels in the northeast Pacific, for example, rising by approximately 3 dB re 1 µPa per decade from 1965 to 2003 (M. A. McDonald, Hildebrand, and Wiggins 2006). Causes of anthropogenic noise primarily comprise shipping and other marine vessel traffic (Haviland-Howell et al. 2007; J. A. Hildebrand 2009; Ross 2005), high-powered sonars (Bernaldo De Quirós et al. 2019), explosive ordnance (Strehse and Maser 2020), pile driving (Bagočius 2015), offshore surveys, drilling, and offshore wind farms (Nedwell and Howell 2004). These noise polluters cover a frequency spectrum between 5 Hz and 200 kHz (NRC 2003) in different intensities and ranges (Fig. 1). The most significant and primary impact of ocean anthropogenic noise derives from shipping traffic (J. Hildebrand 2004; R. I. McDonald, Kareiva, and Forman 2008; Merchant et al. 2012; Ross 2005), affecting almost all marine areas worldwide. It is essential to differentiate between sources of sound associated to vessel traffic. Sound sources include cavitation (Ross 1979), engine noise (Hattori, Nakamachi, and Sanada 1985), hull resonances (Oppenheimer and Dubowsky 2003), flow noise (Wang and Moin 2000) and propeller noise (Tani et al. 2016). The primary acoustic emissions from boats predominantly arise from cavitation, which spans frequencies from 5 Hz to 100 kHz (Ross 1979) and engine noise, where the vessel’s speed is a critical factor, with an increase of 4-6 dB in sound pressure level (SPL) with a doubling of speed (Kipple and Gabriele 2003). Certain hull characteristics can further amplify SPLs (Javier et al. 2023). For example, cargo vessels (173 m length, 16 knots) produce 192 dB re 1 µPa on a bandwidth of 40-100 Hz (J. A. Hildebrand 2009). Larger supertankers can reach peak values of 195 dB re 1µPa at around 30 Hz (NRC 2003), and smaller boats (e.g. rigid inflatable boats with outboard engines) produce sound at a peak range of 1 to 5 kHz with SPLs of 150-180 dB re 1 µPa at 1m (Erbe 2002), Fig. 1). Generally, larger vessels produce higher noise levels than smaller boats, with their frequency emissions more concentrated at lower frequencies. The differences in frequency ranges can largely be attributed to the hull design; longer vessels facilitate slower and more pronounced oscillations, resulting in lower frequency outputs. In contrast, smaller vessels with shorter hulls tend to produce sound at higher frequencies and cover a broader acoustic spectrum (Lu et al. 2013). This highlights the intricate relationship between vessel size, hull characteristics, and the resultant underwater noise profiles. Underwater explosions constitute another significant source of anthropogenic noise in the low frequency range, predominantly arising from ship shock trials and military bomb testing. These activities can generate peak SPLs of up to 272 dB re 1 µPa per pound of Trinitrotoluene (TNT) (NRC 2003; Shin 2004; Urick 1975), (Fig. 1). While underwater construction activities also contribute to marine noise pollution, their impact has been markedly reduced in recent decades, except for impulsive sound production due to pile driving for offshore wind parks (Stöber and Thomsen 2021). Furthermore, military testing has been restricted to specific regions of the oceans to mitigate broader ecological effects. Another contributor to underwater noise pollution is sound navigation and ranging (SONAR). SONAR systems have many applications, including military operations, scientific research, and fisheries (Gonzalez-Socoloske and Olivera-Gómez 2023; Hansen 2013; Hożyń 2021; Wei, Duan, and An 2022). Regardless of their specific use, all SONAR systems, whether military, commercial, or scientific, emit sound frequencies within specific ranges, producing high sound pressure levels necessary for effective operation at significant depths or over extended distances. Depending on their operational parameters, SONAR systems can achieve amplitude levels as high as 250 dB re 1 µPa in low-frequency ranges below 1 kHz up to a few kHz (Fig. 1). In contrast, others operate in higher frequencies, from 100 kHz to 500 kHz, also achieving similar amplitude levels (Bjørnø 2017). Commercial SONAR systems typically function at frequencies up to 1 MHz, generating considerably lower sound pressure levels than their military counterparts (Andrews 2003), Fig. 1).

**Figure 1:**
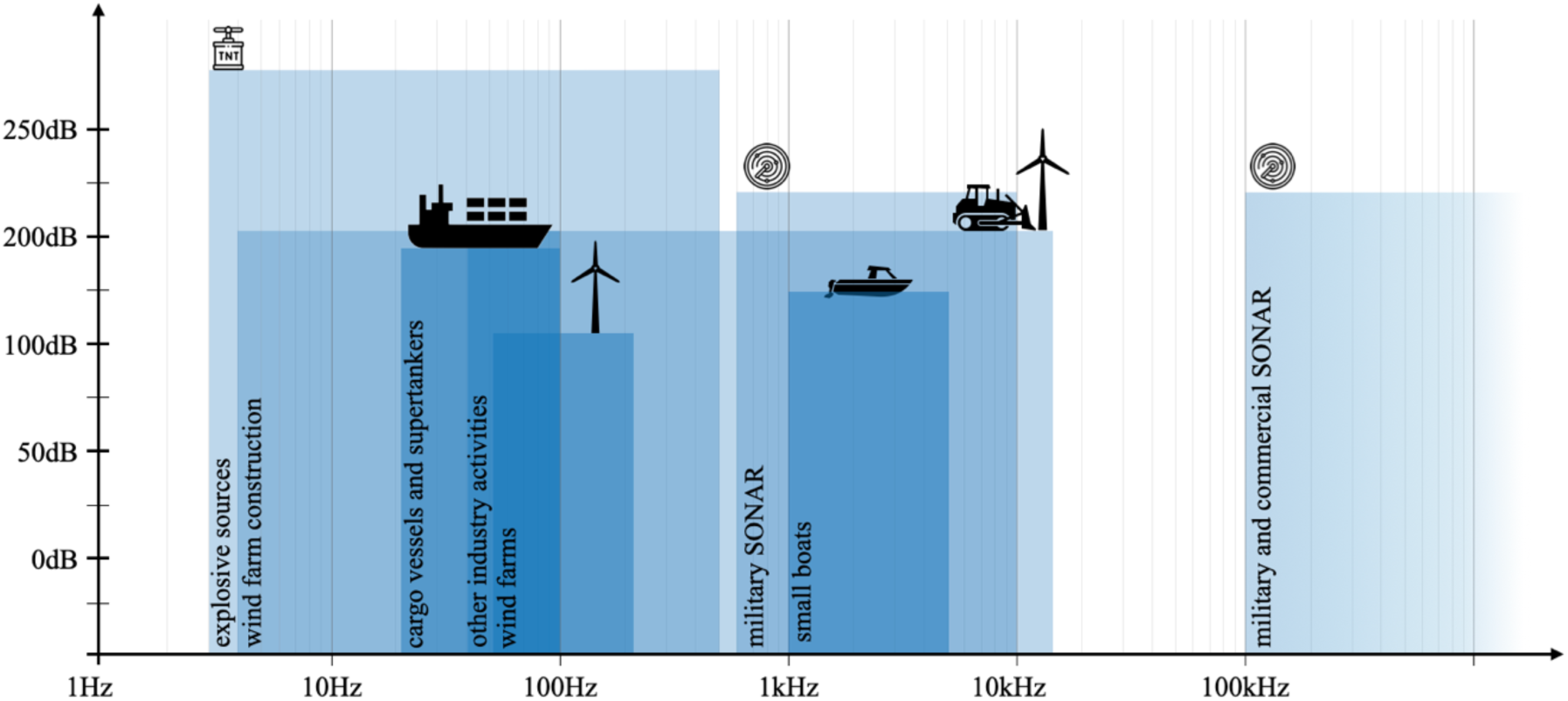
Schematic drawing of anthropogenic noise pollution in the ocean. Frequency ranges (Hz) and peak values (blue areas) of sound pressure levels (dB re 1 µPa) for different sources of anthropogenic sounds. Overlapping opaque blue shows superimpositions of different sound sources in the same spectrum. Sound intensities and spectra originate from peer-reviewed literature. (Andrews 2003; Bagočius 2013; Bernaldo De Quirós et al. 2019; Bjørnø 2017; Chahouri, Elouahmani, and Ouchene 2022; Erbe 2002; Gonzalez-Socoloske and Olivera-Gómez 2023; Hansen 2013; Hattori, Nakamachi, and Sanada 1985; Haviland-Howell et al. 2007; J. Hildebrand 2004; J. A. Hildebrand 2009; Hożyń 2021; Javier et al. 2023; Kipple and Gabriele 2003; Lu et al. 2013; M. A. McDonald, Hildebrand, and Wiggins 2006; R. I. McDonald, Kareiva, and Forman 2008; Merchant et al. 2012; Nedelec et al. 2016; Nedwell and Howell 2004; NRC 2003; Oppenheimer and Dubowsky 2003; Ross 1979, 2005; Shin 2004; Solé et al. 2023; Stöber and Thomsen 2021; Strehse and Maser 2020; Urick 1975; Wang and Moin 2000; Wei, Duan, and An 2022)

While mammals, fish, molluscs, and crustaceans have been well studied in respect to noise pollution, other invertebrates, such as ascidians, remain poorly investigated (Solé et al. 2023). This is largely due to limited knowledge on their sound perception, as well as a lack of data on their behavioural responses to sound. The ascidian *Halocynthia papillosa*, which inhabits coastal waters at a depth of 10 meters or more in the Mediterranean Seas, is exposed to ship noise because of its substrate-attached, sessile lifestyle. Research on ascidian responses to noise is scarce, with studies primarily focusing on model organisms like *Ciona intestinalis*, where changes in settlement, metamorphosis, and survival have been observed (J. I. McDonald et al. 2014). More recently, *Styela plicata* was found to react to boat noise and two other stimuli (White, Edwards, and Ambrosio 2021). A recent study by Varello et al. (2023) presents first research on behaviour responses to ultrasonic stimuli in an ascidian (*S. plicata*) highlighting the need for further research on sound perception and behavioural responses in both natural (field) and controlled (laboratory) settings.

In contrast to airborne or waterborne sound, subsurface vibrations are a separate but equally consequential component of underwater noise pollution. These vibrations propagate through solid materials such as sediments, rocks, and human-made structures, creating mechanical disturbances that can be perceived by sessile marine organisms (Hawkins and Popper 2017; Roberts and Elliott 2017). As ascidians are sessile organisms, they may be susceptible to stimuli due to their direct attachment to the substrate. Whereas motile organisms can actively avoid unfavourable acoustic conditions, ascidians are constantly exposed to the vibrations of the substrate, which requires physiological or behavioural adaptations. It has been shown that mechanical stimulation can induce behavioural changes in ascidians, such as siphon closure, altered feeding activity or withdrawal responses, suggesting a mechanosensory role in the perception of their environment (Burighel et al. 2003; Mackie et al. 2006). Ascidians may rely on the structural properties of their mantle to translate signals into mechanoreceptive cues (Rigon et al. 2013). It has been hypothesised that the sensory structures in the siphonal epithelium and larval papillae function similarly to proprioceptors or vibration-sensitive mechanoreceptors, enabling them to detect micromovements of the substrate (Anselmi et al. 2024; Burighel et al. 2003). In the larval stage, this vibration perception may play a role in the decision to settle, with substrate texture, stability, and movement contributing to habitat selection (Wakai et al. 2021). These vibrational perceptions could provide an evolutionary advantage by ensuring that ascidian larvae settle on stable substrates that minimise future disturbance. In a further context, anthropogenic vibrations generated by shipping traffic, industrial activities, and offshore infrastructure may interfere with ascidians’ sensory perception, leading to disruption of feeding efficiency, settlement patterns, or reproductive success.

In contrast to natural vibration sources such as currents or bioturbation, artificial substrate vibrations can introduce atypical frequency ranges or unpredictable intensities that may lead to stress responses. Understanding the role of substrate vibrations in ascidian ecology is, therefore, crucial for assessing the broader impact of underwater anthropogenic noise. While past studies have focused primarily on waterborne noise effects on fish and marine mammals, increasing attention is being directed toward benthic invertebrates and sessile species (Hawkins and Popper 2017; Roberts and Elliott 2017; Solé et al. 2023). No studies have – to the best of our knowledge – isolated different components of underwater noise (e.g. substrate vibrations and sound pressure waves) with regard to their effect on sessile marine invertebrates. Investigating ascidian vibration thresholds, sensory mechanisms, and long-term adaptation strategies will be essential in evaluating how these sessile filter feeders navigate their dynamic acoustic environment. In this study, we present three experiments with the ascidian *H. papillosa* that a) determine which acoustic frequency ranges induce behavioural alterations, b) decouple response to sound pressure from response to substrate vibrations and determine response threshold for vibration intensities at different frequencies, and c) investigate habituation effects to prolonged substrate vibration exposure.

## 2. Materials and methods

The present study aimed at investigating the perception of and reaction to sound of *Halocynthia papillosa* in its natural habitat and in a controllable laboratory environment. In the first stage, a field experiment was conducted in which the animals were exposed to artificially induced controlled sound signals *in situ*. This was followed by laboratory tests under standardised conditions, allowing the presentation of the same sound stimuli while minimising environmental variability. In the second stage, the components of underwater sound and substrate vibrations were decoupled to determine conclusively whether the observed contractions were triggered by waterborne sound pressure or by structure-borne sound in the substrate. Several individuals were randomly assigned to different stimulus sequences (Table 1) throughout all experimental sections, reducing potential habituation effects. “Sham” trials without sound stimulation or vibration elements were included to distinguish spontaneous contractions from stimulus-induced behaviour. In a third stage, habituation trials were performed to investigate whether repeated exposure to vibration stimuli alters the contraction response over time.

**Table 1:** Overview of experimental stimuli and test conditions. In experiment 1, the sequence of the frequencies played back to the ascidians corresponded to the subdivided groups A and B. In experiment 2, the sequence of the frequencies was randomized.

| Experiment | Frequencies (Hz) | Test type | Sound file characteristics |
| --- | --- | --- | --- |
| 1: Field Trial<br>(Fig. 2a) | A: 600, 200, 1000<br>B: 100, 800, 400, 1500 | Acoustic stimulus<br>presentation, in situ<br>conditions | Acclimatization 10min<br>Four times alternating: saw tooth tone (30sec) and<br>silence interval (9min 30sec) |
| 1: Laboratory Trial<br>(Fig. 2b) | A: 600, 200, 1000<br>B: 100, 800, 400, 1500 | Acoustic stimulus<br>presentation,<br>laboratory conditions | Acclimatization 10min<br>Four times alternating: saw tooth tone (30sec) and<br>silence interval (9min 30sec) |
| 2: Decoupling<br>Substrate Vibration<br>(Fig. 2c) | 50, 80, 100, 200, 300, 400,<br>500, 600, 700, 800, 900,<br>1000, 1500 | Isolated substrate<br>vibration | Sine tones in 2s blocks, amplitude increase/decrease,<br>airtight sealed setup to reduce sound pressure and<br>particle movement. |
| 2: Decoupling<br>Underwater Sound<br>(Fig. 2d) | 80, 100, 200, 300, 400, 500,<br>600, 700, 800, 900, 1000,<br>1500 | Isolated sound pressure<br>without substrate<br>vibration | Sine tones in 2 sec blocks, cover with flexible<br>membrane for induction of sound pressure |
| 3: Habituation Trial<br>(Fig. 2c) | 100 | Habituation to<br>substrate vibrations at<br>1 m/s<sup>2</sup> | Sine tone in 2 sec blocks. Acceleration at 1 m/s<sup>2</sup>. |

### 2.1. Sample collection and study area

For the field experiments (Fig. 2a), *H. papillosa* individuals were kept in their natural habitat. The experiments were performed while SCUBA diving (Pula, Croatia, N44.83°, E13.84°) from 18^th^-21^st^ August, 2022, at 8-18 m depth (T_surface_ = 24.2-25.0°C, T_8-18m_ = 17-23°C, Sal_surface_ = 37–39 PSU). Surface temperature was measured with a digital thermometer (Greisinger GMH2710-I). During the dive, depths and temperatures were monitored with an Apple Watch Ultra (serial number: D706QFXCW0) as a dive computer. Surface salinity was monitored with a handheld refractometer (Aqua Medic). The animals for the laboratory experiment (Fig. 2b) were collected at the same site near Pula on 21^th^ June, 2022 (T_18m_ = 18°C).

**Figure 2:**
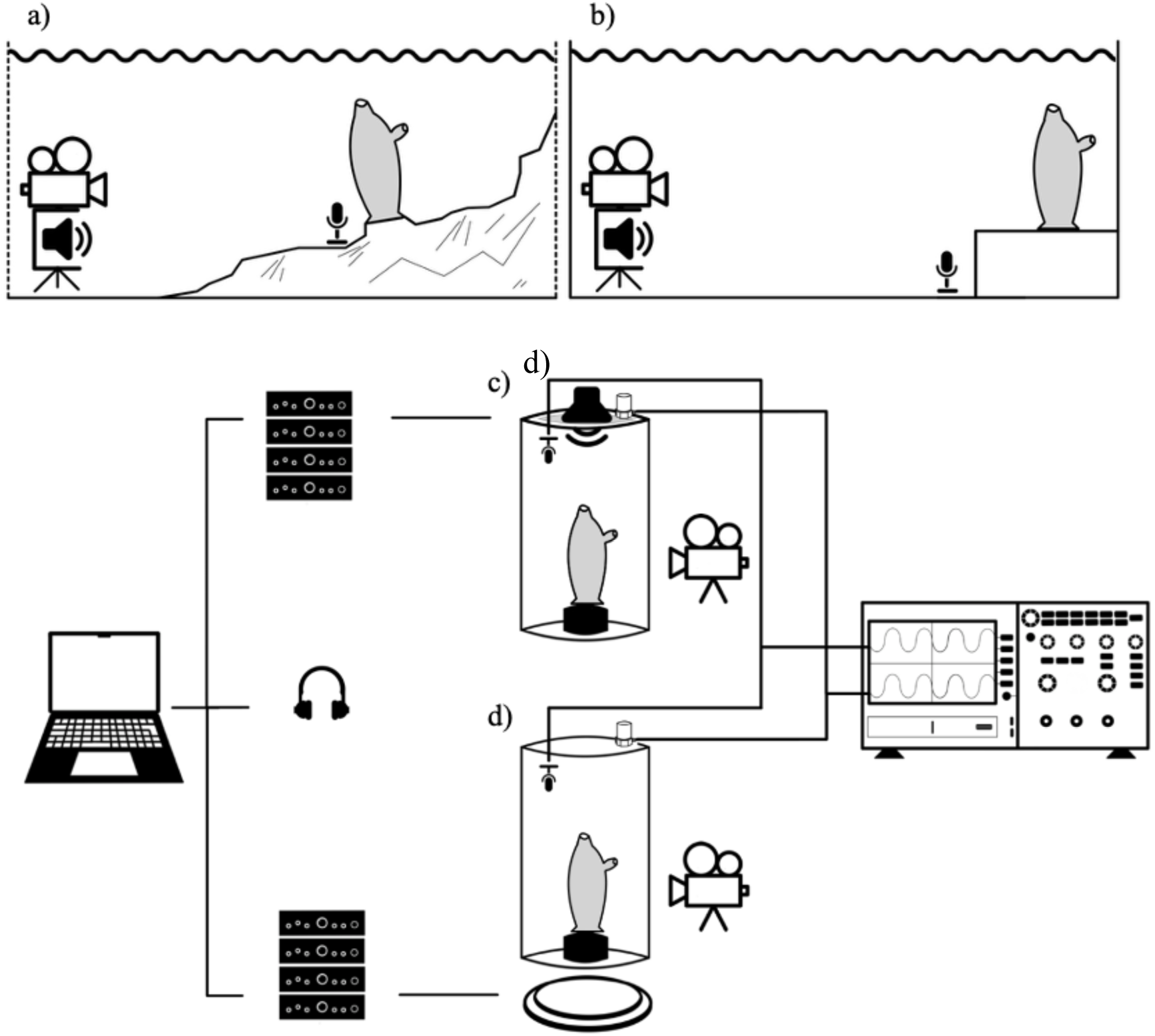
Experimental setup. **a)** Experimental setup in the field experiment (experiment 1), consisting of a self-engineered frame is (left-hand side), containing a waterproof sealed speaker, an attached action camera on the top, and adjustable tripod feet below. The microphone icon illustrates the hydrophone in front of the ascidian. **b)** Laboratory experiment (experiment 1) with the same frame as in the field setup but without tripods. A Eurobox® enclosed the setup in a cooling chamber. **c**,**d)** Setup of the decoupling experiment (experiment 2). A cylinder (outer diameter 40 cm, height 40 cm) with two different lids for the different test conditions was used (rigid lid in **d,** lid with a flexible membrane in **c**). The cylinder was placed on a vibration plate (**d**) or a vibration-free table (**c**). A speaker on top of the membrane (**c**) generated sound pressure in the cylinder and kept vibrations to a minimum. In experiment 3 (habituation), the same setup as in **d** was used.

The animals for the decoupling and habituation test (Fig. 2c) were collected from the island of Elba (42.75758° N, 10.40144° E) from 9^th^ to 13^th^ July, 2024. The temperature at the sampling site on the island of Elba was recorded using an Apple Watch Ultra 2 (serial number: JK9RLLJQXR) as a dive computer (T_surface_ = 22 °C, T_30m_ = 16 °C). Sampling took place at a depth of between 15 and 32 metres. Average salinities of 37.08 PSU were measured with a Hydrolab HL7 multiparameter probe. Animal collection was done utilizing a diving knife, a hammer, and a chisel. Including the adhering limestone to avoid cutting off the roots of the animals. Collected specimens were transported to Ruhr University Bochum (RUB) submersed in individual plastic containers containing seawater from the collection site. Each container accommodated six to ten organisms and an aeration stone. Containers were kept inside a cooling box maintained at a consistent temperature of 16°C. Two water exchanges were performed during the 14-hour travel duration that was approximately the same from Pula and from Elba. In the laboratory, the animals were transferred to a different aquarium (123L) and acclimatized to laboratory conditions on a specially developed, adaptable rack that separates the animals within the aquarium and ensures a predefined space. Each individual was fixed to a movable frag plug using underwater glue. With these frag plugs, animals could later easily be moved to the experimental setups. During the all transfer procedures it was ensured that the animals were not exposed to air. The aquarium system was continuously cooled to 16°C via an external chilling device (TECO TK150) and maintained through a carbon filter (AquaMedic Carbolit 4mm), a plastic ball filter (AquaMedic Miniballs), and a ceramic ring filter (AquaNova NCR-0.5) provided in an external technical tank, complemented by continuous water exchange facilitated by an externally located pump (EHEIM compactON 1000, 15W). Following acclimatization with natural seawater sourced from the sampling site, artificial seawater was prepared using AquaMedic Reef Salt, achieving a salinity of 38-40 PSU. A water exchange of a quarter of the aquarium volume was conducted weekly. A light cycle (12h light/12h dark) was regulated by an Aquarsky fixture (6500K, 21W). Acoustic aquarium insulation was achieved with acoustic foam varying from 2.5cm to 6cm in thickness. Decoupling of vibrations from the aquarium’s incoming and outgoing labels and hoses was accomplished using a rubber mat. Daily feeding of the specimens was conducted, providing a total volume of 20 ml per individual per day (∼1 x 10^7^ c/ml) *Nannochloropsis salina* was delivered during six equally spaced cycles through a feeding pump (EasyGrow).

### 2.2. Field trial on the perception of underwater sound

For this study, we engineered a durable aluminium frame specifically designed for field trials, which effectively accommodated a splash-proof speaker (BLIX Forever, model BS-850) that was sealed in vacuum bags to make it waterproof, and an action camera (LAMAX W7.1). The structure incorporated two adjustable tripod legs to ensure stable positioning on the uneven seabed, with the speaker and camera maintained at a distance of 30-40 cm from the selected individuals (Fig. 2a). To facilitate subsequent automated evaluations based on grey value analyses, solitary standing individuals in front of a light background, and an atrial siphon oriented to the left, were explicitly chosen for inclusion in the experiment. For half of the measurements, a hydrophone (HydroMoth®) was placed in front of the animal to check whether the frequencies played were correctly transmitted into the water and thus reached the animal accurately. Dive durations extended up to one hour, during which multiple setups (two to four) were deployed and measured by different divers. Care was taken to maintain a sufficient spatial separation between setups to mitigate potential interference of the multiple setups. Following the sonication phase, the distance between animal and the setup, water temperature, and depth were recorded, and the frame was subsequently disassembled. Each individual was tested only once during the sonication process to uphold the integrity of the data collected. The sonication procedure lasted 40 minutes, comprising a 10-minute acclimatization phase followed by the presentation of four distinct frequencies within one group, each lasting for 30 seconds, interspersed with silent intervals of 9 minutes and 30 seconds (Table 1).

### 2.3. Sound protocol and data processing

Specified sinusoidal and sawtooth frequencies, ranging from 100 Hz to 1500 Hz in predefined intervals (Table 1), were generated using Audacity (Version 3.5). The acclimatization, sonication, and rest phases were standardized in advance and adjusted according to the requirements of the respective trials. Sound profiles (A, B or silence) varied within the trials, however, MP4 video recordings were made for all experiments with either a LAMAX W7.1 or a GoPro Hero 8 black. The video data were synchronized with the hydrophone data and set to 1fps to reduce the memory size for post-processing. Videos were manually revised and checked for environmental disturbances such as animals, divers, or boat noise interactions, and acquired data with environmental disturbances were skipped in the analysis. After loading the videos into FIJI (ImageJ, Version 2.16.0) as hyperstacks, the colormap was set to greyscale, and the threshold was adjusted to enhance the ratio of the animal to the background. For the numerical recording of the contraction, masks of the maximum occupying 2D space of the animal during the whole trial were created by a maximum or minimum intensity projection (MIP). The mask’s mean grey value for every frame (resp. 1 sec) was extracted to a table for further analysis. Mean grey values represented the relative expansion of the individual with high means representing a large expansion and low means a strong contraction (black animal on white background). Mean grey values were standardized by the largest possible grey value an individual showed as this represented the full expansion of the animal. This, therefore, resulted in a proportional expansion of each individual at each time point with 1 being equal to a full expansion. To quantify the contraction that occurred at the time of onset of an acoustic signal, integrals were calculated over 30 seconds directly prior to the acoustic signal and over 30 seconds after the signal. Integral ratios were then calculated as the 30 sec integral after the signal, divided by the 30 sec integral prior to the signal. This resulted in one integral ratio per signal type (different frequencies) per individual.

### 2.4. Laboratory test on the perception of underwater sound

To better control for variations in environmental parameters and eliminate potential interference with environmental influences, tests similar to the experiments in the field were carried out in a cooling chamber at RUB. The animals were placed individually in an Eurocontainer® (60 × 40 × 40 cm) of which inside walls and ground were covered with sound and vibration insulation mats. A GoPro Hero 8 black recorded the animal behaviour using 1080p and 24 fps with linear lens settings. The temperature in the room was constant between 14°C and 16°C, making an external cooling unit and filter system unnecessary for the short duration of the experiment (1 hour), thus preventing noise and flow disturbances. The same acoustic stimuli (Table 1) and speaker (BS-850) used in the field trial were used. A stronger vacuum was applied to the vacuum bag sealed speaker to mimic pressure underwater. A sawtooth tone of different frequencies was presented at ten-minute intervals with a distance similar to the field trials. Because of the handling of the animals, the acclimatization time was set to 15 minutes. The following procedure and the number of sonication cycles followed the same scheme as in the field, using the same protocol, sound, and video recording. Image postprocessing also took place in Fiji and integral ratios were calculated as for the field experiment.

### 2.5. Decoupling substrate vibration and underwater sound

A special test and measurement system was developed to determine whether the contractions observed in *H. papillosa* are triggered by sound pressure waves in the water or by vibrations in the substrate. The central component was a vibrating plate with an upper and a lower stainless-steel plate, which could be excited via electrodynamic shakers (Visaton EX 60 S - 8 Ω). An additional speaker (Visaton 8Ω 8.5cm 10W) generated targeted acoustic signals, making switching between the two stimulation components possible. A transparent, cylindrical container made of Plexiglas^®^ (outer diameter 40 cm, height 40 cm, wall thickness 0.5 cm, lid/bottom 2.5 cm each) served as a test chamber. The animals could be secured firmly to the ground by fitting the frag plug inside a matching holder. For substrate vibration tests, the cylinder was completely filled with water. The cylinder lid was sealed airtight to prevent water movement and minimise the sound pressure waves (Fig. 2d). For the second trial, an alternative lid design with an integrated elastic membrane was manufactured to enable the introduction of sound pressure over the membrane lid while reducing substrate vibrations (Fig. 2c). The cylinder was mounted on a vibration-free table to remove external uncontrollable vibrations. The hydrophone (MS10CM) and accelerometer (MMF KD37) enabled the simultaneous recording of sound pressure and vibration amplitude, with all measurements conducted using an oscilloscope (Tektronix TDS 1001B). Calibration of the hydrophone took place in an anechoic chamber with a reference sound source, while the accelerometer signal was calibrated with a Polytec LASER Doppler vibrometer (VibroGO VGO-200). The stimuli (frequencies between 50 Hz and 1500 Hz, Table 1) were applied in two-second blocks, the amplitude of which increased and decreased to the desired strength within the two seconds.

### 2.6. Behavioural reaction analysis to stimulus exposure

The contraction behaviour of the oral siphon was used as the response parameter, as *H. papillosa* reacts to even minimal stimulation with a visible contraction retraction. A central aspect of the study was determining the threshold at which a response to a stimulus occurs. For this purpose, the selected frequencies and accelerations of the vibrations were tested in randomized order. Each tested acceleration was repeated 30 times per frequency. The observations were carried out in a blind test, in which the recording of whether or not a contraction of the oral siphon occurred was made. Of the 30 stimuli presented, 15 were actual stimuli, while 15 were “sham” stimuli and served as control. After the placement into the test cylinder, an acclimatization period of 30 minutes was provided. In each session, the individuals were exposed to a maximum of six stimuli consisting of a randomised mixture of real and sham stimuli. Between each stimulus, there was an acclimatization period of 10 minutes. After each session of six repetitions, the animals were returned to the holding aquarium and given a rest period of 48 hours before being tested again. These vibration experiments identified a range in which the threshold for stimulus perception lies. The experimental setup was adapted for sound perception using a similar procedure. Here, the technically highest possible amplitude was used, and the vibration acceleration was reduced as much as possible as a side effect to test whether the animals reacted to the high sound pressure or the substrate vibrations.

### 2.7. Habituation test

To test habituation behaviour to repeating presented stimuli, 12 individuals were exposed 25 times with the same vibration stimulus (Table 1) 100 Hz, 2 s duration, 1 m/s^2^ in a laboratory test. The acoustic stimulation was started after a 30-minute acclimatization phase followed by alternating stimuli exposure (2 s) and silent interval (10 min). This resulted in a cycle duration of two hours with nine stimuli continuously played without interaction, after which the water in the cylinder was exchanged. After two hours, the animal was changed and used again the next day. The BORIS software (Version 9.0.0) was used to measure temporal parameters to evaluate the behavioural data. In this regard, two behavioural parameters were measured: firstly, the latency time until contraction after the start of the stimulus. Secondly, the time it took the animal to relax fully after the beginning of the contraction. In addition, photos were analysed in Datinf^®^ Measure, on which the projected area of the sea squirt could be marked. In this way, the changes in expansion could be determined, especially during the transition from maximum expansion to contraction. The recorded behavioural data was used to assess whether the reaction latencies increased or the contraction strengths decreased - both indicators of habituation. To check on possible habituation, the reaction intensity after the first stimulus was compared to the reaction intensity after the last stimulus. To ensure that a longer stay in the experimental set-up did not influence the behaviour of the animals, the behaviour of six animals without vibration stimulus was observed in the same setting at the same time points without exposure to stimuli. No contraction was observed.

### 2.8. Statistical analysis

All statistical analyses and figures were generated using RStudio (Version 2024.12.1+563). To compare contraction behaviour between laboratory and field conditions in experiment 1, a linear mixed model was applied using the lme4 package (Bates et al. 2015), with the different sound stimuli (100 Hz, 200 Hz, 400 Hz, 600 Hz, 800 Hz, 1000 Hz and 1500 Hz) and place (laboratory and field) as fixed factors, and the individuals as random factor. Significance was determined using t-tests based on Satterthwaite’s method. Frequencies were compared against a ratio of 1.0, which was added as a reference data point for each individual, as 1 represented no change in body size from before to after the stimulus (no contraction). The interaction term for the fixed factors was included and kept. To identify possible reaction thresholds in experiment 2, chi-square tests were performed, comparing the frequency of contractions at all acceleration and sound pressure levels to establish the threshold at which significant contraction occurred during vibration and SPL exposure. Pearson correlation analyses were used to examine whether substrate vibration (acceleration) increases were associated with elevated sound pressure (dB). In experiment 3, time courses of contraction and relaxation times in response to repeated stimuli were analysed by linear regression. A Wilcoxon signed-rank test was performed to compare contraction intensity—defined as the percentage reduction in body area—between the first and last stimulus.

## 3. Results

### 3.1. Ascidian reaction to sound in the field and laboratory

In the first experiment, the reaction of *H. papillosa* during exposure to sound stimuli of different frequencies were tested in the field and in the laboratory. The video recordings showed that all tested frequencies could provoke a contraction response in the ascidians. However, for the higher frequencies (1000 Hz and 1500 Hz) , and especially in the laboratory, contractions were rare. To distinguish between randomly occurring contractions and sound-induced contractions, we tested for the effect of sound on the quantifiable expansion of the animal. Relative expansion of directly after the onset of a stimuli was quantified as integral ratio (expansion of an animal during 30 seconds after the stimuli, divided by the expansion during 30 seconds before the stimuli). Relative expansion significantly decreased upon exposure to sounds of 100 Hz, 200 Hz, 400 Hz, 600 Hz and 800 Hz (REML, Table 2, Fig. 3). Overall, relative expansion did not differ between animals exposed in the laboratory and in the field. However, a significant interaction between frequency and treatment location was found at 400 Hz and 600 Hz, and a marginally insignificant interaction at 800 Hz and 1000 Hz. Field-treated animals reacted stronger to these frequencies than lab-treated animals (Fig. 3). To account for potential confounding factors, vibrations generated as a secondary effect of the loudspeaker were measured retrospectively in the laboratory setup. For technical limitations, however, this parameter could not be determined in the field setup. The measured amplitudes of the vibrations corresponding to the tested frequencies ranged from 0.471 m/s^2^ at 100 Hz to 0.149 m/s^2^ at 1500 Hz, with an approximately linear decline in values up to 600 Hz, followed by fluctuations at higher frequencies (Table 3). The lowest amplitude was observed at 1000 Hz (0.067 m/s^2^).

**Figure 3:**
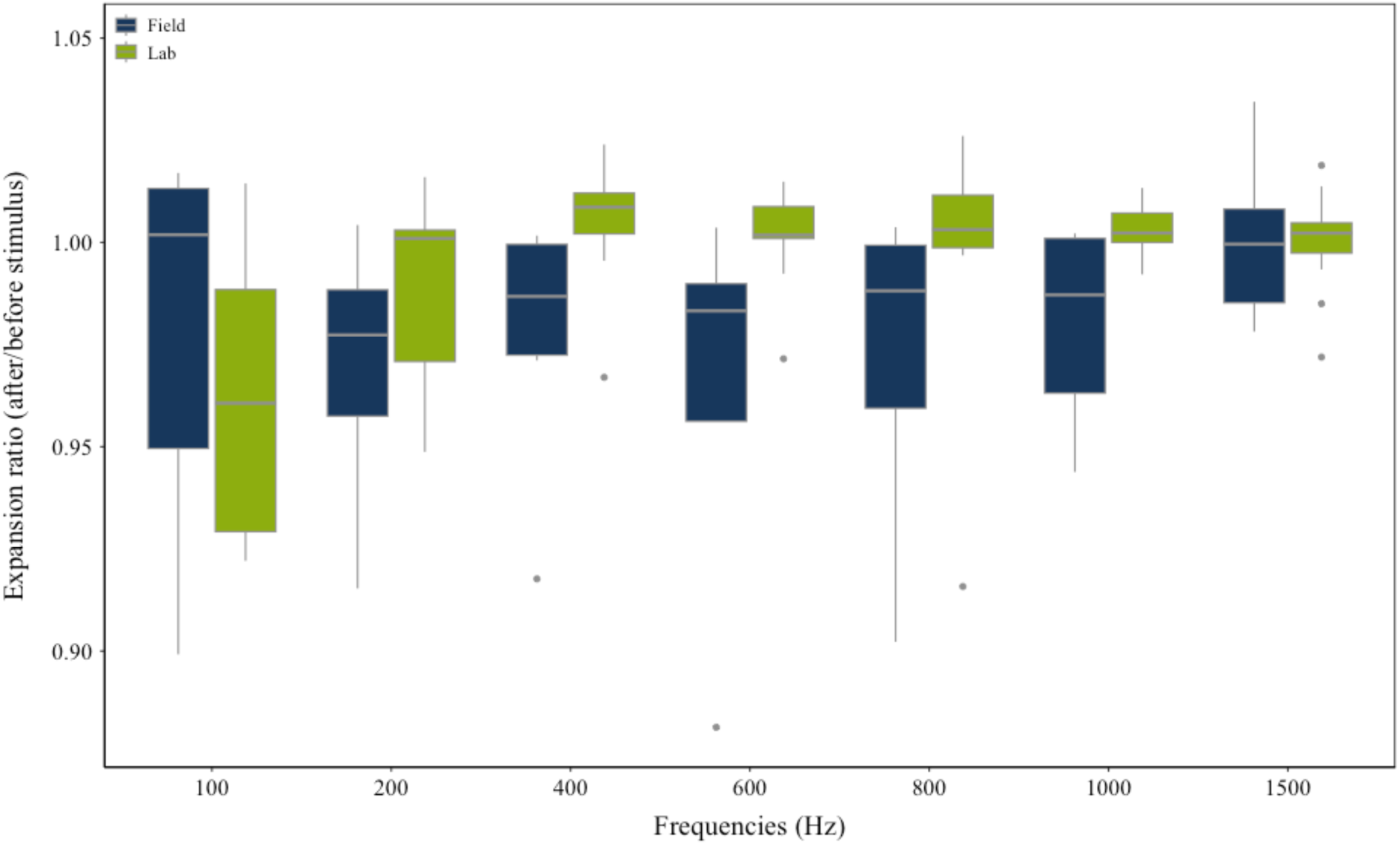
Relative expansion of *Halocynthia papillosa* in response to sound stimuli of different frequencies in the laboratory and field (experiment 1). Expansion ratios quantify body posture changes following stimulus onset (ratio > 1 = expansion, < 1 = contraction, = 1 = no change). The ratio was calculated by dividing the integrated surface area covered by the animal in the 30 s after stimulus onset by the area in the 30 s before. Statistical tests were conducted against a null hypothesis of no response (ratio = 1). Significance values are reported in Table 2.

**Table 2:** Output of the mixed model (lme4 package in R) assessing place-independent effects of exposure to sound stimuli of different frequencies (100-1500 Hz), and the interaction between place (laboratory and field) and the different sound stimuli. Place and frequency were treated as fixed factors and individuals as random factor (Model formula: Integral ratio ∼ Frequency*Place + (1|Individual)). All frequencies were compared against a ratio of 1, Pairwise comparisons were made using Satterthwaite’s method (lmerModLmerTest). Significance codes: (•) p < 0.1, (*) p < 0.05, (**) p < 0.01, (***) p < 0.001. REML criterion at convergence = -634.5, median of scaled residuals = 0.0354, n_(observations)_ = 156, n_(individuals)_ = 28

| Condition | Effect size | SE | df | t-value | P-value | Significance niveau |
| --- | --- | --- | --- | --- | --- | --- |
| Intercept | 1.000 | 0.006 | 127.6 | 159.862 | < 0.001 | *** |
| 100 Hz | -0.021 | 0.011 | 129.9 | -2.006 | 0.047 | * |
| 200 Hz | -0.032 | 0.012 | 136.3 | -2.604 | 0.010 | * |
| 400 Hz | -0.021 | 0.010 | 128.3 | -2.074 | 0.040 | * |
| 600 Hz | -0.038 | 0.012 | 136.3 | -3.063 | 0.003 | ** |
| 800 Hz | -0.027 | 0.012 | 132.1 | -2.236 | 0.027 | * |
| 1000 Hz | -0.021 | 0.014 | 134.8 | -1.801 | 0.074 | . |
| 1500 Hz | 0.000 | 0.010 | 126.2 | 0.014 | 0.989 |  |
| Lab | -0.000 | 0.008 | 112.5 | -0.002 | 0.998 |  |
| 100 Hz – Lab | -0.016 | 0.013 | 126.7 | -1.225 | 0.223 |  |
| 200 Hz – Lab | 0.022 | 0.015 | 132.4 | 1.465 | 0.145 |  |
| 400 Hz – Lab | 0.026 | 0.012 | 124.0 | 2.135 | 0.035 | * |
| 600 Hz – Lab | 0.040 | 0.015 | 132.4 | 2.754 | 0.006 | ** |
| 800 Hz – Lab | 0.027 | 0.015 | 128.8 | 1.852 | 0.066 | .. |
| 1000 Hz – Lab | 0.023 | 0.014 | 130.1 | 1.721 | 0.088 | . |
| 1500 Hz - Lab | 0.000 | 0.012 | 123.1 | 0.037 | 0.970 |  |

**Table 3:** Amplitudes of vibrations corresponding to the different frequencies produced by the setup of the laboratory trials in experiment 1.

| Frequency (Hz) | Vibration amplitude (m/s<sup>2</sup>) |
| --- | --- |
| 100 | 0.471 |
| 200 | 0.420 |
| 300 | 0.353 |
| 400 | 0.236 |
| 500 | 0.167 |
| 600 | 0.084 |
| 700 | 0.094 |
| 800 | 0.077 |
| 900 | 0.118 |
| 1000 | 0.067 |
| 1500 | 0.149 |

### 3.2. Reaction thresholds to substrate vibration

To identify the lower limit at which *H. papillosa* contracts in response to vibrations in experiment 2, vibration amplitudes (acceleration in m/s^2^) were randomly varied in small steps around a previously assumed response threshold. This threshold was derived by comparing accelerations at which the animals showed a significant contraction (Chi-square test) to the next lower acceleration that did not trigger a measurable response. For technical reasons, this range varies from 0.3-1.0 m/s^2^ at different frequencies (grey area, Fig. 4). Repeated tests (n=30) were performed for each step to account for variations in individual response thresholds. The animals contracted significantly when the vibration amplitudes exceeded a specific threshold value, which varied depending on the frequency.

**Figure 4:**
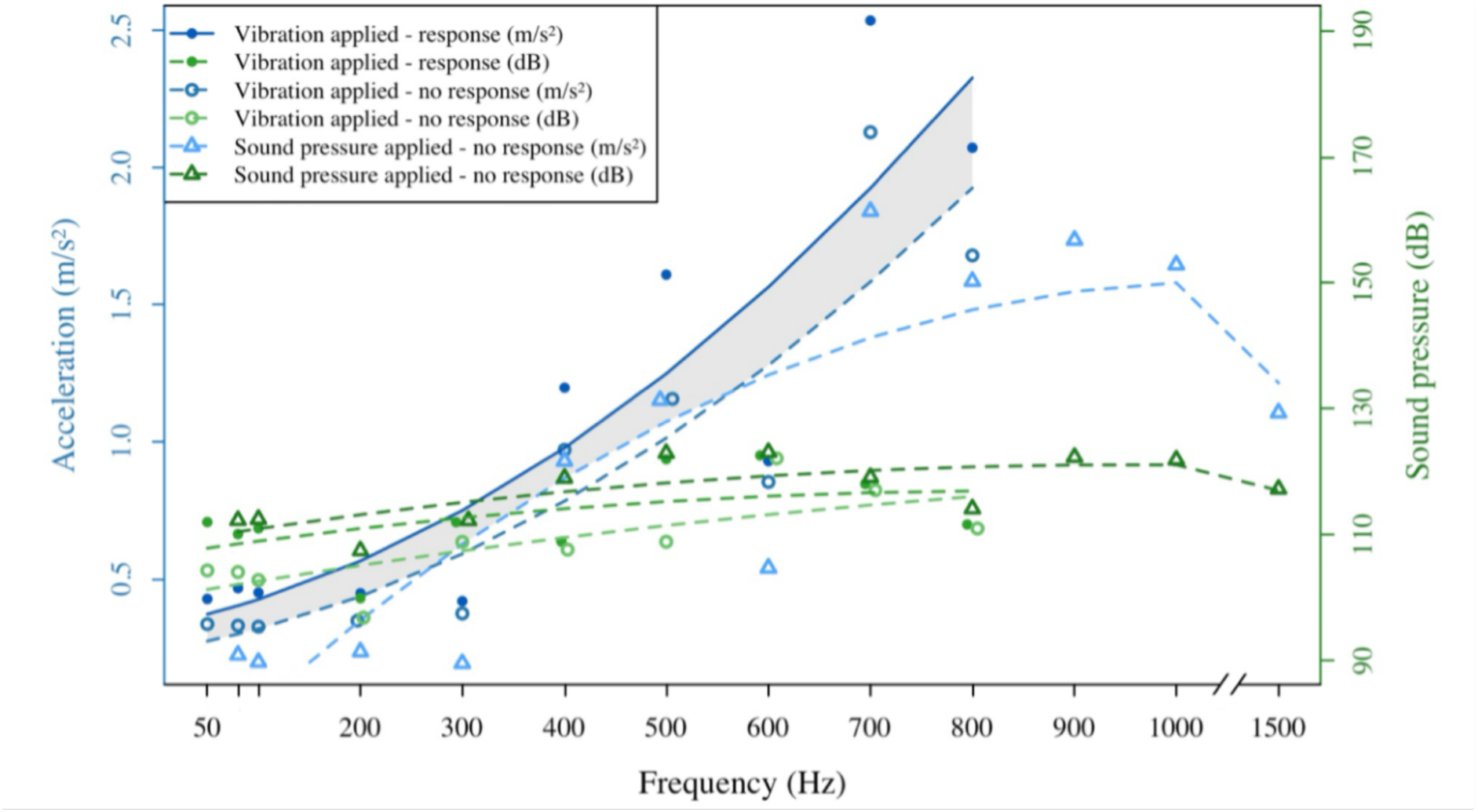
Acceleration and sound pressure over the frequencies for different treatments. The blue data points and lines show the parameters for measured vibrations (left y-axis, in m/s^2^). Green data points and lines show the parameters for measured sound pressure waves (right y-axis, in dB re 1 µPa). Filled markers and the solid line show a significant behavioural response of the animals, while no response was measured for unfilled markers and dashed lines. The grey area marks the area of the vibration acceleration in which the actual reaction threshold of the ascidians is located. In order to test whether the measured behavioural response of the animals was due to the vibration or the sound pressure, the experimental setup was adapted. This experiment is represented by triangles, showing no response of the animals at higher sound pressure levels (dark green), with reduced substrate vibrations (light blue).

In contrast, no significant behavioural response occurred at the next lower acceleration, mainly about 0.1 m/s^2^ below the higher acceleration condition. Chi-square tests confirmed that the ascidians had a clear threshold beyond which the probability of contraction increased sharply (Table 4) (p < 0.001 for the difference between amplitude ranges below the threshold and above the threshold). The threshold varied between the individual frequencies, with exposures to 50 - 600 Hz causing contractions at lower amplitudes (Fig. 4). Higher frequencies (>800 Hz) provoked no significant response.

**Table 4:** Mean acceleration (m/s^2^) and mean sound pressure level (Ø dB) under different frequency conditions. Statistical significance is indicated by p-values, with the corresponding Cramér’s V-values and chi-square statistics reflecting the strength and significance of the correlations. The measurements are given for different accelerations of the vibration plate, with p-values < 0.05 indicating statistically significant differences.

| Condition (Hz) | Mean acceleration<br>($\text{m/s}^2$) | Mean SPL<br>(dB re 1 $\mu\text{Pa}$) | p-value | Cramér's V | Chi-square value |
| --- | --- | --- | --- | --- | --- |
| 50 | 0.332 | 104.3 | 0.065 | 0.047 | 3.394 |
| 50 | 0.424 | 111.9 | 0.001 | 0.006 | 10.80 |
| 80 | 0.327 | 104.0 | 0.065 | 0.047 | 3.394 |
| 80 | 0.463 | 110.0 | 0.003 | 0.011 | 8.571 |
| 100 | 0.323 | 102.7 | 0.058 | 0.044 | 3.589 |
| 100 | 0.447 | 110.9 | 0.046 | 0.039 | 3.968 |
| 200 | 0.345 | 96.8 | 0.136 | 0.067 | 2.222 |
| 200 | 0.445 | 99.9 | 0.046 | 0.039 | 3.968 |
| 300 | 0.371 | 108.8 | 0.121 | 0.064 | 2.400 |
| 300 | 0.416 | 111.9 | 0.002 | 0.008 | 9.600 |
| 400 | 0.968 | 107.6 | 0.065 | 0.053 | 3.407 |
| 400 | 1.195 | 108.8 | 0.005 | 0.015 | 7.886 |
| 500 | 1.154 | 108.8 | 0.121 | 0.064 | 2.400 |
| 500 | 1.608 | 121.9 | 0.002 | 0.008 | 9.600 |
| 600 | 0.850 | 122.0 | 0.142 | 0.069 | 2.160 |
| 600 | 0.927 | 122.5 | 0.002 | 0.008 | 9.600 |
| 700 | 2.128 | 117.0 | 0.068 | 0.048 | 3.333 |
| 700 | 2.536 | 118.0 | 0.046 | 0.039 | 3.968 |
| 800 | 1.678 | 111.0 | 0.195 | 0.081 | 1.677 |
| 800 | 2.071 | 111.5 | 0.046 | 0.039 | 3.968 |
| 900 | 3.145 | 120.7 | 0.666 | 0.149 | 0.186 |
| 1000 | 2.181 | 120.0 | 0.283 | 0.097 | 1.154 |
| 1500 | 1.362 | 97.4 | 0.142 | 0.069 | 2.160 |

**Table 5:** Mean acceleration (m/s^2^) and mean sound pressure level (dB) under different frequency conditions in the adapted test set-up with higher sound pressure and lower acceleration. Statistical significance is indicated by p-values, with corresponding Cramér’s V-values and chi-square statistics reflecting the strength and significance of the correlations. Measurements are given for the matched higher sound pressure levels of the vibration plate, with p-values < 0.05 indicating statistically significant differences.

| Condition (Hz) | Mean sound pressure level | Mean acceleration |  |  |  |
| --- | --- | --- | --- | --- | --- |
| | (dB re 1 $\mu\text{Pa}$) | ($\text{m/s}^2$) | p-value | Cramér's V | Chi-square value |
| 80 | 112.2 | 0.220 | 0.099 | 0.057 | 2.727 |
| 100 | 112.4 | 0.193 | 0.142 | 0.069 | 2.160 |
| 200 | 107.4 | 0.232 | 0.121 | 0.064 | 2.400 |
| 300 | 112.2 | 0.189 | 0.121 | 0.064 | 2.400 |
| 400 | 118.9 | 0.927 | 0.705 | 0.153 | 0.144 |
| 500 | 122.8 | 1.149 | 0.143 | 0.069 | 2.143 |
| 600 | 123.0 | 0.537 | 0.068 | 0.048 | 3.333 |
| 700 | 119.0 | 1.839 | 0.283 | 0.097 | 1.154 |
| 800 | 114.0 | 1.584 | 0.283 | 0.097 | 1.154 |
| 900 | 122.2 | 1.734 | 0.283 | 0.097 | 1.154 |
| 1000 | 121.8 | 1.644 | 0.143 | 0.069 | 2.143 |
| 15000 | 117.2 | 1.104 | 0.361 | 0.110 | 0.833 |

### 3.3. Reaction thresholds to sound pressure waves

The reaction of *H. papillosa* to increased sound pressure with substrate vibrations kept at a minimum was tested using a cylinder lid with an elastic membrane that reduced the direct mechanical coupling of the cylinder and cylinder base, keeping the measured acceleration below the determined threshold. Various sine tones (Table 1) were played at volumes identical to or above the side effects measured during the substrate vibration test. During the experiment the acceleration did not rise above the previously determined threshold values. Despite high sound levels (up to 130 dB re 1 µPa), no significant contractions occurred if the vibration amplitude was lower than the threshold (Chi-square tests, p > 0.05 in all frequency ranges, n = 30).

### 3.4. Correlation between substrate vibration and sound pressure

Pearson’s correlation analyses were conducted for all experimental conditions in experiment 2 to investigate the relationship between substrate vibration (acceleration) and the associated sound pressure. This analysis aimed to clarify whether proportional increases in sound pressure consistently accompany increases in acceleration and whether this relationship might explain the observed behavioural responses. In the above-threshold condition, where significant behavioural responses were triggered, no significant correlation was observed between acceleration and sound pressure (r = 0.457, p = 0.1838). Conversely, the below-threshold condition, where no significant behavioural responses were observed, exhibited a stronger correlation between acceleration and sound pressure (r = 0.625, p = 0.0531). This correlation was marginally significant, indicating that while the animals did not contract, the physical coupling between substrate vibration and sound pressure was more consistent at these lower amplitudes. The test condition with higher sound pressure, without substrate vibrations, showed the strongest correlation (r = 0.656, p = 0.0148), emphasising a robust linear relationship between acceleration and acoustic pressure. However, despite the high sound pressure levels (up to 130 dB re 1 µPa), no significant behavioural responses were observed when the substrate vibrations remained below the contraction threshold.

### 3.5. Habituation test: Contraction and relaxation time

The contraction duration was measured as the time from the onset of the stimulus to the peak contraction of the ascidian (Fig. 5 a). This metric provides insights into the speed of the organism’s behavioural response to vibratory stimulation. Repeated exposure to the stimulus significantly decreased contraction duration (linear regression: p < 0.001, R^2^ = 0.60, n = 17). This progressive reduction suggests a habituation effect, where the initial arousal response diminishes over time. The relaxation time, defined as the interval from peak contraction to full recovery of the sea squirt, was also significantly reduced over repeated stimulations (Fig. 5 b; p < 0.001, R^2^ = 0.64, n = 17).

**Figure 5:**
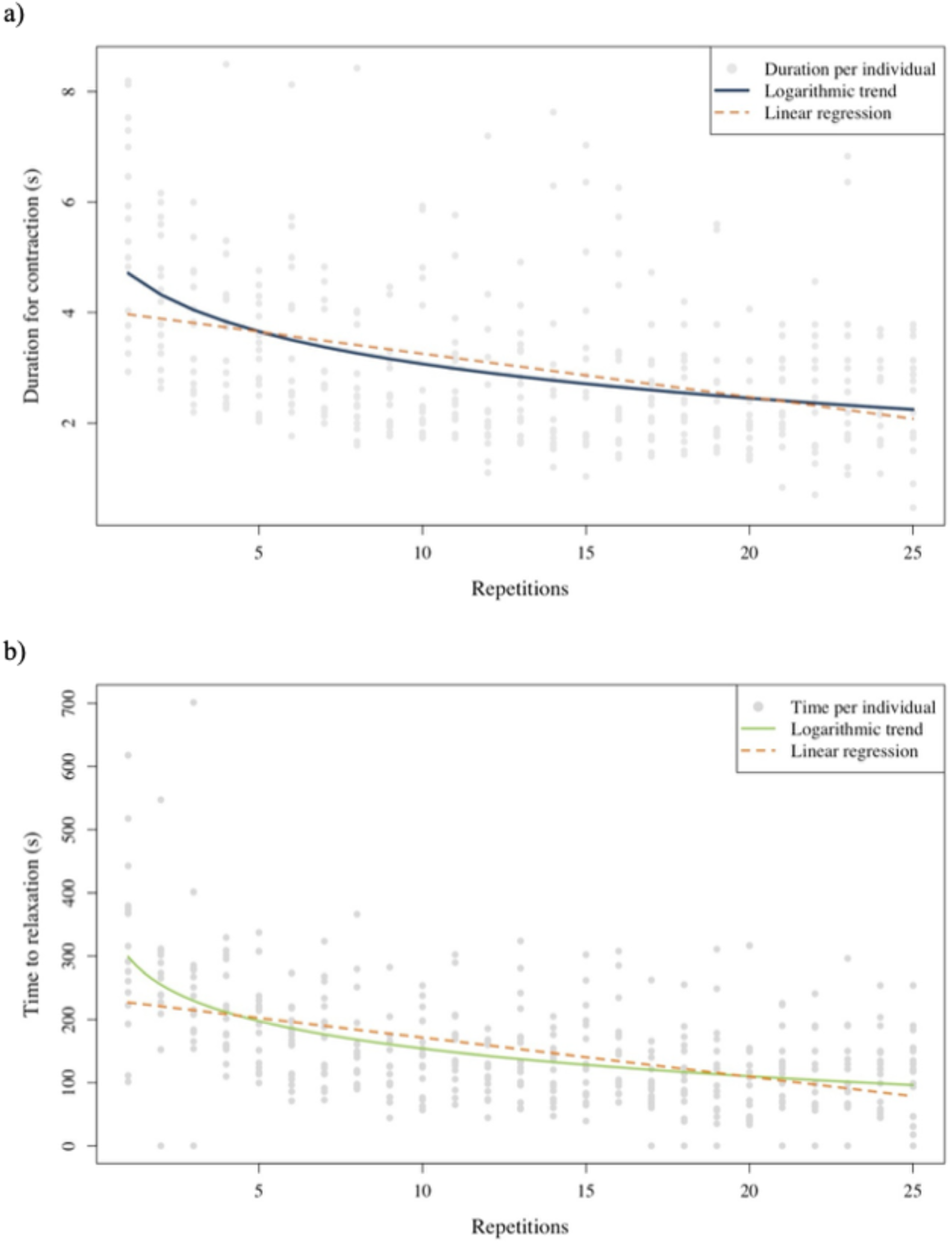
Changes in contraction duration (**a**) and relaxation time (**b**) in *Halocynthia papillosa* over 25 stimulus repetitions (vibrations of 100 Hz, 1 m/s^2^). Grey markers represent time measurements per individual, blue and green solid lines show the locally estimated scatterplot smoothing trends (logarithmic), and the orange dashed lines represent a linear regression model. Both contraction duration and relaxation time decreased significantly with repeated stimulation (p < 0.001 for contraction and relaxation). N = 17.

**Fig. 6:**
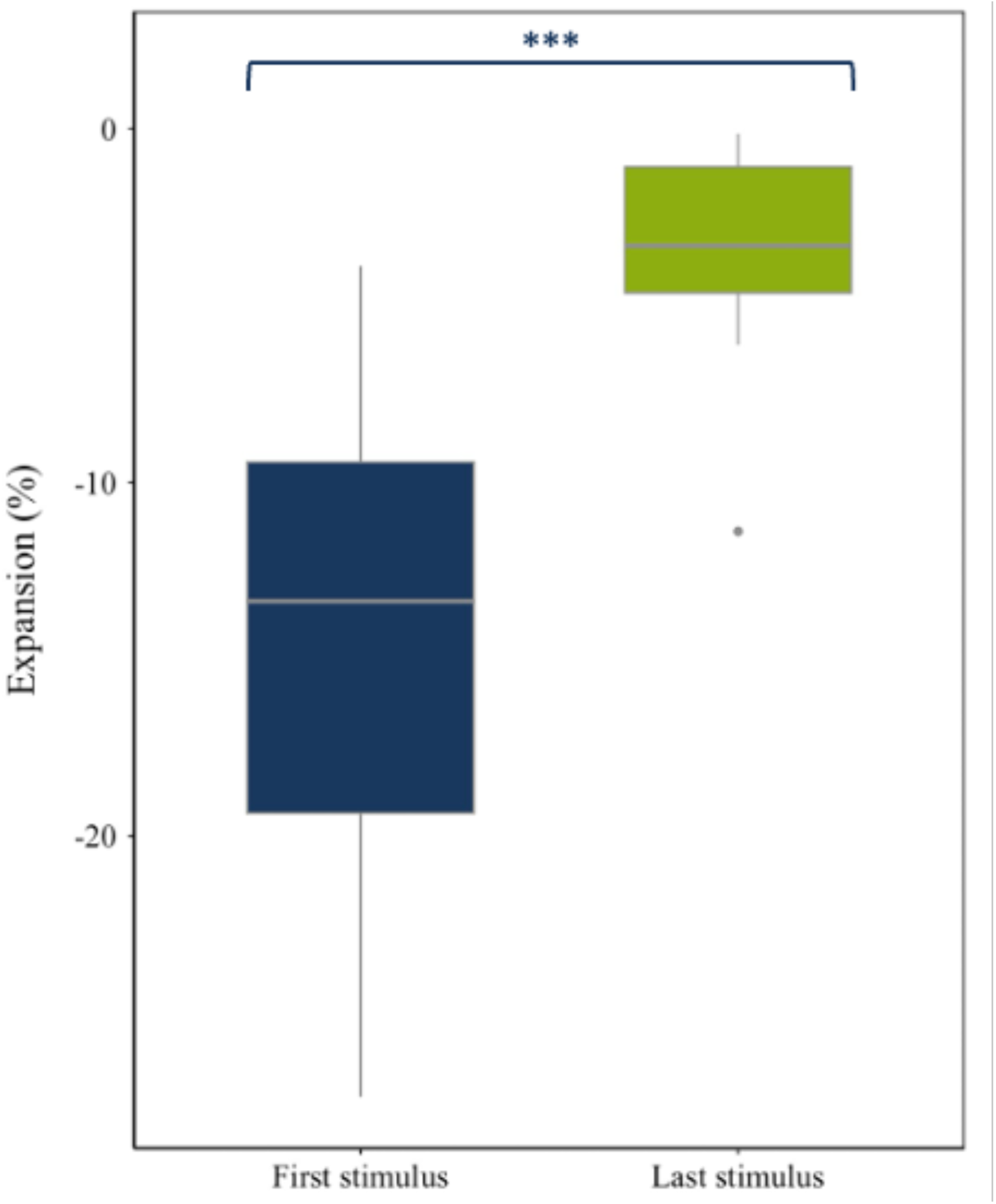
Change (median ± 1.5 * IQR) in expansion (%) of *Halocynthia papillosa* following the first and last of 25 stimuli (vibrations of 100 Hz, 1 m/s^2^) in the habituation experiment. N=17. ***P < 0.001.

### 3.6. Habituation test: Contraction intensity

Contraction intensity was analysed to further assess habituation by quantifying the projected body area (expansion) of an animal after the first and last (25^th^) stimulus. A highly significant reduction in contraction rate was observed between the first and last stimulus (Wilcoxon signed-rank test, V = 0, p < 0.001, Cliff’s Delta = 0.92, n = 17), indicating a strong habituation effect. The contraction rate decreased consistently over repeated trials, following a logarithmic decay pattern characteristic of habituation. Initially, subjects exhibited contraction intensities averaging 14.1 % ± 6.6 % of their resting area, which declined to 3.3 % ± 2.8 % after the last stimulus (M ± SD).

## 4. Discussion

Our results demonstrate that *Halocynthia papillosa* contracts in response to acoustic stimuli of a spectrum between 50 and at least 800 Hz. By decoupling water-transmitted sound pressure waves and substrate vibrations we could furthermore show that *H. papillosa* rather responds to the substrate vibrations caused by acoustic stimuli, than to the sound pressure directly. Moreover, observed habituation effects suggest that the ascidians possess plasticity in their sensory-motor responses, modulating their contraction behaviour when repeatedly exposed to vibrational stimuli. We discuss the implications of these findings in the context of sensory ecology, behavioural plasticity, and environmental impact while also acknowledging the study’s limitations and future research directions.

This study shows that contraction behaviour in *H. papillosa* is exhibited in response to rather low sound frequencies (100 Hz - 800 Hz), whereas no strong behavioural responses occurred at frequencies above 800 Hz. The pronounced difference in contraction response to frequencies above 400 Hz between field- and laboratory-exposed animals suggests that perception sensitivity is highest at very low frequencies (200 Hz or lower). The fact that in the laboratory setup (a box lined with acoustic foam to reduce sound reflection and vibrations) contraction responses were less expressed above 200 Hz (Fig. 3) than in the field where the animals were attached to harder and more rigid substrates, hints that secondarily caused substrate vibrations compose the main stimulus to provoke contraction. This is supported by the vibration intensity measurements of the lab setup, which showed an almost linear decrease in vibration perception from 100 Hz to 600 Hz, followed by consistently low vibration intensities at higher frequencies. Overall, the decreasing contraction response with increasing frequencies we observed suggests a threshold effect, where *H. papillosa* is most sensitive to lower-frequency substrate vibrations. This can be due to the biological relevance and abundance of low frequency vibrations, as this frequency range is commonly produced by benthic invertebrates, including crabs, during locomotion and feeding (Patek and Caldwell 2006; Roberts and Breithaupt 2016). Our study is the first study to demonstrate which frequency ranges in substrate vibrations can be perceived by ascidians. Past studies have either focused on sound in general (White, Edwards, and Ambrosio 2021) or simply correlated the abundance of biofouling ascidians to the proximity to a vibrating generator on a vessel hull (McDonald et al. 2014). None of these studies looked at the vibration quality and intensity. One study (Varello et al. 2023) suggested that the ascidians *Ciona intestinalis*, *Ascidiella aspersa* and *Styela plicata* perceive vibrations in the ultrasonic range (30-35 kHz). At first sight, our findings contradict the conclusion of this study as we demonstrate that the sensitivity to vibrations decreases at frequencies higher than 800 Hz in *H. papillosa*. It is unlikely, that different species of solitary ascidians that occupy similar habitats would differ to such a large extend in their ability to perceive sound signals. We suggest that the ascidians in the study conducted by Varello et al. (2023) reacted to secondary vibrations produced by the sonication devices inside the polystyrene box that was used for the experiments. Vibrations were not measured in the study, but sound pressure waves were measured with a hydrophone (HydroMoth) and were concluded to produce peaks at >30 kHz. The low cost HydroMoth, however, has been tested to produce recordings comparable to high-tech devices up to 15 kHz and produces high background noise at higher frequencies (Lamont et al. 2022). It, therefore, is not an adequate device to draw conclusion on ultrasound. The power spectral densities produced by the sonication devices, which were reported by Varello et al. (2023) show power peaks in the low frequencies. Therefore, we suggest that the ascidians in that study did not react to ultrasound but to vibrations caused by the sound pressure waves inside the box that were in the lower frequency range (<1000 Hz). This highlights the importance of measuring substrate vibrations and sound pressure waves in experiments investigating underwater acoustics, especially when conducted in relatively small containers or tanks in the laboratory. Our results also show that the dual experimental approach in marine bioacoustics (replicating in the field and in the laboratory) should be conducted whenever possible. Field trials are important for testing the ecological relevance of acoustic stimuli while laboratory trials allow a controlled mechanistic analysis.

The assumption on sensitivity to substrate vibrations drawn from the experiment replicated in the field and laboratory was further analysed in a controlled experimental approach. This led to a key finding of this study, which is that *H. papillosa* does not show any contraction response to increased sound pressure levels when substrate vibrations are minimised. No response to sound pressure was observed when vibrations were underneath frequency-specific thresholds, even at high sound pressure levels of over 130 dB re 1 µPa. Our study systematically eliminated substrate vibrations during high-intensity sound pressure exposure, ensuring that sound pressure was the only stimulus affecting *H. papillosa*. The absence of contraction responses under these controlled conditions provides compelling evidence that this species does not detect or respond to sound pressure alone, supporting the notion that ascidians rely solely on mechanoreception via the substrate or particle motion in the water body. These novel results suggest that ascidians do not have specialised auditory structures for the perception of water-born sound pressure and respond primarily to direct mechanical disturbances of the substrate. This response may represent an adaptive mechanism to mitigate predation risk. Previous studies have demonstrated that ascidians are predated upon by benthic organisms, particularly crabs, which mechanically manipulate or crush them to access internal tissues (Bernárdez, Freire, and González-Gurriarán 2000). Crabs generate low-frequency substrate vibrations during locomotion and feeding, and the ability of ascidians to detect such mechanical stimuli may confer a selective advantage by enabling contraction in response to approaching threats (Roberts and Breithaupt 2016). Furthermore, ascidians frequently occur in high-density aggregations where mechanically induced conspecific contractions may propagate as substrate-borne disturbances, potentially eliciting coordinated group responses (Petersen and Svane 1995). This suggests that vibratory cues play a fundamental role in both individual and collective behavioural adaptations in ascidians. The failure to adequately differentiate between acoustic pressure waves and substrate vibrations in previous studies has likely resulted in an overestimation of the auditory capabilities of sessile marine invertebrates. While behavioural modifications in response to noise exposure have been widely documented, few studies have controlled for the possibility that these responses were driven by substrate vibrations rather than by sound pressure. The prevailing assumption that marine invertebrates primarily respond to sound pressure has profoundly influenced discussions on anthropogenic noise pollution. Most studies investigating the effects of underwater noise—particularly those examining shipping noise, seismic surveys, and offshore construction—have focused predominantly on sound pressure levels, assuming that this parameter alone accounts for observed behavioural or physiological changes (Hawkins and Popper 2017; Nedelec et al. 2016; Popper and Hawkins 2018). Similarly, large-scale reviews addressing the ecological consequences of anthropogenic noise have emphasized sound pressure while largely neglecting vibratory components (Solé et al. 2023). This methodological bias has shaped both experimental designs and regulatory frameworks, reinforcing the notion that marine invertebrates primarily perceive and respond to auditory stimuli, despite mounting evidence suggesting that substrate-borne vibrations are ecologically more relevant (Roberts and Elliott 2017; Roberts and Howard 2022).

To date, investigations focusing explicitly on vibratory sensitivity in marine invertebrates remain scarce. Only a limited number of taxa—specifically four arthropod and four molluscan species—have been subjected to controlled experimental analyses regarding their responses to substrate-borne vibrations (Ellers 1995; Roberts and Howard 2022). Notably, for sessile filter feeders outside of Bivalvia, such as ascidians, no such studies have been conducted to date. The present study challenges this paradigm, demonstrating that vibratory stimuli, rather than sound pressure, constitute the principal sensory input for sessile benthic invertebrates such as ascidians. These findings underscore the necessity of refining experimental approaches and regulatory policies to account for substrate-borne vibrations when assessing the impacts of underwater noise on marine ecosystems. While much research on underwater noise has focused on marine mammals and fish, sessile invertebrates like *H. papillosa* may be more affected by vibrations than waterborne sound pressure. The implications of this distinction are significant, particularly in the context of anthropogenic noise impact assessments. If sessile marine invertebrates primarily respond to substrate vibrations rather than waterborne sound pressure, the focus of conservation efforts and regulatory measures should shift accordingly. Noise mitigation strategies that solely reduce airborne or waterborne sound pressure, such as modifications in ship propeller design or noise barriers, may be insufficient in protecting benthic communities from disturbance. Instead, measures that minimize substrate vibrations, such as altering construction techniques or establishing buffer zones around high-vibration areas, could reduce environmental stress on sessile organisms more effectively. Many anthropogenic activities, such as shipping, pile driving, and offshore energy installations, generate significant substrate vibrations in the frequency range of 100–600 Hz, precisely where *H. papillosa* exhibited its strongest contraction responses (Burighel et al. 2003; Hawkins and Popper 2017; Olivier et al. 2023; Roberts and Elliott 2017). Previous studies on other benthic organisms have demonstrated that prolonged exposure to substrate vibrations can disrupt physiological functions, impair larval settlement, and alter community structures (Joo and Kim 2024; Roberts et al. 2015; Salmon and Atsaides 1969).

The perception of sound and vibrations will depend on the substrate a sessile organism lives attached to. Sound waves can generate secondary vibrations in solid substrates, the strength of which depends on material, porosity, and density (Elias, Mason, and Hebets 2010; Rodríguez and Desjonquères 2019). While rigid substrates such as rock or metal efficiently convert sound into vibrations, softer sediments absorb this energy and attenuate signal transmission (Stritih and Čokl 2012). Our study used a homogeneous, rigid substrate (stainless steel plate) to generate vibrations specifically. However, in natural habitats, different substrates could significantly influence the perception of sounds by *H. papillosa*. Rigid substrates could elicit stronger vibrations and, thus, more intense responses, while the same sounds could be less biologically relevant in sediments (Gordon and Uetz 2011). A systematic database on substrate-dependent sound vibration coupling is still lacking. This makes it difficult to assess which noise sources are particularly relevant for benthic organisms. For example, ship noise could trigger stronger vibration responses in rocky coastal areas than in sandy regions. A targeted investigation of these effects could explain why earlier studies show contradictory results on the reaction of benthic invertebrates to noise (Hebets et al. 2008; Hill 2009).

The habituation response to repeated vibration stimuli we observed in *H. papillosa* is consistent with classical models of non-associative learning, in which repeated exposure to a non-noxious stimulus leads to a diminished response over time (Thompson and Spencer 1966). The latency of the contraction decreased, the relaxation time shortened, and the intensity of the contraction decreased significantly on repeated trials. This suggests that *H. papillosa* can modulate its defensive responses to optimise energy expenditure and avoid unnecessary contraction behaviour in response to non-threatening stimuli. Such behavioural plasticity is well documented in other sessile marine invertebrates, including bivalves and echinoderms, which also show habituation to mechanical perturbations (Dehaudt et al. 2019; Freas and Cheng 2022). From an ecological perspective, habituation allows ascidians to discriminate between persistent background noise and acute disturbance, ensuring that their contraction reflex remains an adaptive defence mechanism and does not result in unnecessary energy loss.

## 5. Conclusion

This study provides compelling evidence that *H. papillosa* responds to substrate vibrations rather than sound pressure waves, highlighting the importance of mechanoreception in sessile marine invertebrates. By systematically decoupling these two stimuli, we demonstrated that contraction responses are most pronounced at low frequencies (≤ 200 Hz) and diminish at frequencies above 800 Hz, suggesting a biologically relevant sensitivity range below 200 Hz. Additionally, the observed habituation effects indicate that *H. papillosa* exhibits behavioural plasticity, allowing it to modulate its responses to repeated vibrational stimuli. From an ecological perspective, our findings emphasize the need to reconsider the impact of anthropogenic noise pollution on benthic organisms, shifting the focus from waterborne sound pressure to substrate vibrations. Many human activities, including shipping, offshore construction, and energy extraction, generate vibrations within the frequency range that triggers contraction responses in *H. papillosa*. Understanding how prolonged exposure affects individual fitness and population dynamics will be crucial for conservation efforts. Future research should explore the long-term effects of chronic substrate vibrations, the neurophysiological mechanisms underlying vibrational perception, and comparative studies across different sessile invertebrates. Expanding our knowledge of how marine organisms perceive and respond to their acoustic environment will improve noise mitigation strategies and ensure the protection of benthic communities in increasingly noise-polluted marine ecosystems.

## CRedit authorship contribution statement

**Til Böttner:** Conceptualization, Methodology, Investigation, Data curation, Formal analysis, Validation, Visualization, Writing – original draft, Writing reviewing & editing. **Lukas Hessel:** Methodology, Investigation, Data curation, Formal analysis, Validation, Visualization, Writing – original draft, Writing reviewing & editing. **René Ortmann:** Investigation, Formal analysis, Writing reviewing & editing. **Wolfgang H. Kirchner:** Conceptualization, Methodology, Writing – original draft, Writing reviewing & editing. **Stefan Herlitze:** Funding acquisition, Conceptualization, Writing – original draft, Writing reviewing & editing. **Mareike Huhn:** Conceptualization, Methodology, Investigation, Data curation, Validation, Formal analysis, Visualization, Supervision, Writing – original draft, Writing reviewing & editing.

## Acknowledgements

We thank the technical workshop of the Faculty of Biology and Biotechnology at Ruhr University Bochum, under the direction of Henning Knoop, for their valuable support in constructing the experimental setups. Thanks to Johanna Wiedling and Diving Pula for their support during the field work.

## Data availability

The data is available at supplementary material 1.

## Ethical approval

All experiments were conducted in accordance with institutional, national, and international guidelines. The study involved marine invertebrates (ascidians) that are not subject to animal welfare regulations. Therefore, no ethical approval was required.

## Declaration of generative AI in scientific writing

AI-assisted technologies (ChatGPT Version 4o, Grammarly) were used in the writing process to improve the readability and language of the manuscript.

## References

Andrews, D.R., 2003. Ultrasonics and acoustics. Encyclopedia of Physical Science and Technology, 17, 269–287. 10.1016/B0-12-227410-5/00800-0

Anselmi, C., Fuller, G.K., Stolfi, A., Groves, A.K., Manni, L., 2024. Sensory cells in tunicates: Insights into mechanoreceptor evolution. Front. Cell Dev. Biol., 12, 1359207. 10.3389/fcell.2024.1359207

Bagočius, D., 2013. Underwater noise level in Klaipėda Strait, Lithuania. Baltica, 26(1), 45–50. 10.5200/baltica.2013.26.05

Bagočius, D., 2015. Piling underwater noise impact on migrating salmon fish during Lithuanian LNG terminal construction (Curonian Lagoon, Eastern Baltic Sea coast). Mar. Pollut. Bull., 92, 45–51. 10.1016/j.marpolbul.2015.01.002

Bates, D., Mächler, M., Bolker, B., Walker, S., 2015. Fitting linear mixed-effects models using lme4. J. Stat. Softw., 67(1), 1–48. 10.18637/jss.v067.i01

Bernaldo De Quirós, Y., Fernandez, A., Baird, R.W., Brownell, R.L., Aguilar De Soto, N., Allen, D., Arbelo, M., et al., 2019. Advances in research on the impacts of anti-submarine sonar on beaked whales. Proc. R. Soc. B, 286, 20182533. 10.1098/rspb.2018.2533

Bernárdez, C., Freire, J., González-Gurriarán, E., 2000. Feeding of the spider crab *Maja squinado* in rocky subtidal areas of the Ría de Arousa (NW Spain). J. Mar. Biol. Assoc. U.K., 80, 95–102. 10.1017/S0025315499001605

Bjørnø, L., 2017. Underwater acoustic measurements and their applications. In: Bjørnø, L. (Ed.), Applied Underwater Acoustics. Elsevier, pp. 889–947. 10.1016/B978-0-12-811240-3.00014-X

Burighel, P., Lane, N.J., Gasparini, F., Tiozzo, S., Zaniolo, G., Candia Carnevali, M.D., Manni, L., 2003. Novel, secondary sensory cell organ in ascidians: In search of the ancestor of the vertebrate lateral line. J. Comp. Neurol., 461, 236–249. 10.1002/cne.10666

Chahouri, A., Elouahmani, N., Ouchene, H., 2022. Recent progress in marine noise pollution: A thorough review. Chemosphere, 291, 132983. 10.1016/j.chemosphere.2021.132983

Dehaudt, B., Nguyen, M., Vadlamudi, A., Blumstein, D.T., 2019. Giant clams discriminate threats along a risk gradient and display varying habituation rates to different stimuli. Ethology, 125, 392–398. 10.1111/eth.12863

Elias, D.O., Mason, A.C., Hebets, E.A., 2010. A signal-substrate match in the substrate-borne component of a multimodal courtship display. Curr. Zool., 56, 370–378. 10.1093/czoolo/56.3.370

Ellers, O., 1995. Discrimination among wave-generated sounds by a swash-riding clam. Biol. Bull., 189, [keine Seitenangabe vorhanden]

Erbe, C., 2002. Underwater noise of whale-watching boats and potential effects on killer whales (*Orcinus orca*), based on an acoustic impact model. Mar. Mamm. Sci., 18, 394–418. 10.1111/j.1748-7692.2002.tb01045.x

Freas, C.A., Cheng, K., 2022. Neuroecology beyond the brain: Learning in Echinodermata. Learn. Behav., 50, 20–36. 10.3758/s13420-021-00492-3

Gonzalez-Socoloske, D., Olivera-Gómez, L.D., 2023. Seeing in the dark: A review of the use of side-scan sonar to detect and study manatees, with an emphasis on Latin America. Lat. Am. J. Aquat. Mamm., 18, 114–124. 10.5597/lajam00301

Gordon, S.D., Uetz, G.W., 2011. Multimodal communication of wolf spiders on different substrates: Evidence for behavioural plasticity. Anim. Behav., 81, 367–375. 10.1016/j.anbehav.2010.11.003

Hansen, R.E., 2013. Synthetic aperture sonar technology review. Mar. Technol. Soc. J., 47(5), 117–127. 10.4031/MTSJ.47.5.5

Hattori, K., Nakamachi, K., Sanada, M., 1985. Prediction of underwater sound radiated from ship’s hull by using statistical energy analysis. [Kein Journal, bitte prüfen]

Haviland-Howell, G., Frankel, A.S., Powell, C.M., Bocconcelli, A., Herman, R.L., Sayigh, L.S., 2007. Recreational boating traffic: A chronic source of anthropogenic noise in the Wilmington, North Carolina Intracoastal Waterway. J. Acoust. Soc. Am., 122, 151–160. 10.1121/1.2717766

Hawkins, A.D., Popper, A.N., 2017. A sound approach to assessing the impact of underwater noise on marine fishes and invertebrates. ICES J. Mar. Sci., 74, 635–651. 10.1093/icesjms/fsw205

Hebets, E.A., Elias, D.O., Mason, A.C., Miller, G.L., Stratton, G.E., 2008. Substrate-dependent signalling success in the wolf spider, *Schizocosa retrorsa*. Anim. Behav., 75, 605–615. 10.1016/j.anbehav.2007.06.021

Hildebrand, J., 2004. Sources of anthropogenic sound in the marine environment. Report to the Marine Mammal Commission. [Bitte prüfen: ggf. graue Literatur, keine Journalangabe]

Hildebrand, J.A., 2009. Anthropogenic and natural sources of ambient noise in the ocean. Mar. Ecol. Prog. Ser., 395, 5–20. 10.3354/meps08353

Hill, P.S.M., 2009. How do animals use substrate-borne vibrations as an information source? Naturwissenschaften, 96, 1355–1371. 10.1007/s00114-009-0588-8

Hożyń, S., 2021. A review of underwater mine detection and classification in sonar imagery. Electronics, 10, 2943. 10.3390/electronics10232943

Javier, R.F., Jaime, R., Pedro, P., Jesus, C., Enrique, S., 2023. Analysis of the underwater radiated noise generated by hull vibrations of the ships. Sensors, 23, 1035. 10.3390/s23021035

Joo, S., Kim, T., 2024. The effect of anthropogenic substrate-borne vibrations on locomotion of the fiddler crab *Austruca lactea*. Mar. Pollut. Bull., 200, 116107. 10.1016/j.marpolbul.2024.116107

Kipple, B.M., Gabriele, C.M., 2003. Glacier Bay underwater noise. Naval Surface Warfare Center Report, [Online]. Available: https://scholar.google.com/scholar_lookup?title=Glacier%20Bay%20Underwater%20Noise [Accessed 24 March 2025]

Lamont, T.A.C., Chapuis, L., Williams, B., Dines, S., Gridley, T., Frainer, G., Fearey, J., et al., 2022. HydroMoth: Testing a prototype low-cost acoustic recorder for aquatic environments. *Remote Sens*. Ecol. Conserv., 8, 362–378. 10.1002/rse2.249

Lu, W., Ling, Y., Song, A., Zeng, H., Ding, W., Xu, B., Gu, S., 2013. Measuring tape-like sampling arm and drill for sampling lunar regolith. Int. J. Adv. Robot. Syst., 10. 10.5772/56361

Mackie, G.O., Burighel, P., Caicci, F., Manni, L., 2006. Innervation of ascidian siphons and their responses to stimulation. Can. J. Zool., 84, 1146–1162. 10.1139/z06-106

McDonald, J.I., Wilkens, S.L., Stanley, J.A., Jeffs, A.G., 2014. Vessel generator noise as a settlement cue for marine biofouling species. Biofouling, 30, 741–749. 10.1080/08927014.2014.919630

McDonald, M.A., Hildebrand, J.A., Wiggins, S.M., 2006. Increases in deep ocean ambient noise in the Northeast Pacific west of San Nicolas Island, California. J. Acoust. Soc. Am., 120, 711–718. 10.1121/1.2216565

McDonald, R.I., Kareiva, P., Forman, R.T.T., 2008. The implications of current and future urbanization for global protected areas and biodiversity conservation. Biol. Conserv., 141, 1695–1703. 10.1016/j.biocon.2008.04.025

Merchant, N.D., Blondel, P., Dakin, D.T., Dorocicz, J., 2012. Measuring acoustic habitats. J. Acoust. Soc. Am., 132, 3394–3404. 10.1121/1.4754429

Nedelec, S.L., Campbell, J., Radford, A.N., Simpson, S.D., Merchant, N.D., 2016. Particle motion: The missing link in underwater acoustic ecology. Methods Ecol. Evol., 7, 836–842. 10.1111/2041-210x.12544

Nedwell, J.R., Howell, D., 2004. A review of offshore windfarm related underwater noise sources. Subacoustech Report, [Online]. Available: www.subacoustech.com [Accessed 24 March 2025]

NRC (National Research Council), 2003. Ocean Noise and Marine Mammals. National Academies Press, Washington, DC. 10.17226/10564

Olivier, F., Gigot, M., Mathias, D., Jezequel, Y., Meziane, T., L’Her, C., Chauvaud, L., Bonnel, J., 2023. Assessing the impacts of anthropogenic sounds on early stages of benthic invertebrates: The “Larvosonic System”. Limnol. Oceanogr. Methods, 21, 53–68. 10.1002/lom3.10527

Oppenheimer, C.H., Dubowsky, S., 2003. A methodology for predicting impact-induced acoustic noise in machine systems. J. Sound Vib., 266, 1025–1051. 10.1016/s0022-460x(02)01450-5

Patek, S.N., Caldwell, R.L., 2006. The stomatopod rumble: Low frequency sound production in *Hemisquilla californiensis*. Mar. Freshw. Behav. Physiol., 39, 99–111. 10.1080/10236240600563289

Petersen, J.K., Svane, I., 1995. Larval dispersal in the ascidian *Ciona intestinalis* (L.): Evidence for a closed population. J. Exp. Mar. Biol. Ecol., 186, 89–102. 10.1016/0022-0981(94)00157-9

Popper, A.N., Hawkins, A.D., 2018. The importance of particle motion to fishes and invertebrates. J. Acoust. Soc. Am., 143, 470–488. 10.1121/1.5021594

Rigon, F., Stach, T., Caicci, F., Gasparini, F., Burighel, P., Manni, L., 2013. Evolutionary diversification of secondary mechanoreceptor cells in Tunicata. BMC Evol. Biol., 13, 112. 10.1186/1471-2148-13-112

Roberts, L., Breithaupt, T., 2016. Sensitivity of crustaceans to substrate-borne vibration. In: Popper, A.N., Hawkins, A.D. (Eds.), The Effects of Noise on Aquatic Life II, Springer, New York, pp. 925–931. 10.1007/978-1-4939-2981-8_114

Roberts, L., Cheesman, S., Breithaupt, T., Elliott, M., 2015. Sensitivity of the mussel *Mytilus edulis* to substrate-borne vibration in relation to anthropogenically generated noise. Mar. Ecol. Prog. Ser., 538, 185–195. 10.3354/meps11468

Roberts, L., Elliott, M., 2017. Good or bad vibrations? Impacts of anthropogenic vibration on the marine epibenthos. Sci. Total Environ., 595, 255–268. 10.1016/j.scitotenv.2017.03.117

Roberts, L., Howard, D.R., 2022. Substrate-borne vibrational noise in the Anthropocene: From land to sea. In: Gordon, T.A.C., Radford, A.N. (Eds.), The Effects of Noise on Aquatic Life III, Springer, Cham, pp. 123–155. 10.1007/978-3-030-97419-0_6

Rodríguez, R.L., Desjonquères, C., 2019. Vibrational signals: Sounds transmitted through solids. Encyclopedia of Animal Behavior, 1, 508–517. 10.1016/B978-0-12-809633-8.90702-7

Ross, D., 1979. Mechanics of Underwater Noise. Pergamon Press, New York. [Online]. Available: https://books.google.de [Accessed 24 March 2025]

Ross, D., 2005. Ship sources of ambient noise. IEEE J. Ocean. Eng., 30, 257–261. 10.1109/JOE.2005.850879

Salmon, M., Atsaides, S.P., 1969. Sensitivity to substrate vibration in the fiddler crab, *Uca pugilator* Bosc. Anim. Behav., 17, 68–76. 10.1016/0003-3472(69)90114-6

Shin, Y.S., 2004. Ship shock modeling and simulation for far-field underwater explosion. Comput. Struct., 82, 2211–2219. 10.1016/j.compstruc.2004.03.075

Solé, M., Kaifu, K., Mooney, T.A., Nedelec, S.L., Olivier, F., Radford, A.N., Vazzana, M., et al., 2023. Marine invertebrates and noise. Front. Mar. Sci., 10, 1129057. 10.3389/fmars.2023.1129057

Stöber, U., Thomsen, F., 2021. How could operational underwater sound from future offshore wind turbines impact marine life? J. Acoust. Soc. Am., 149, 1791–1795. 10.1121/10.0003760

Strehse, J.S., Maser, E., 2020. Marine bivalves as bioindicators for environmental pollutants with focus on dumped munitions in the sea: A review. Mar. Pollut. Bull., 158, 105006. 10.1016/j.marpolbul.2020.105006

Stritih, N., Čokl, A., 2012. Mating behaviour and vibratory signalling in non-hearing cave crickets reflect primitive communication of Ensifera. PLoS ONE, 7, e47646. 10.1371/journal.pone.0047646

Tani, G., Viviani, M., Hallander, J., Johansson, T., Rizzuto, E., 2016. Propeller underwater radiated noise: A comparison between model scale measurements in two different facilities and full scale measurements. Appl. Ocean Res., 56, 48–66. 10.1016/j.apor.2016.01.007

Thompson, R.F., Spencer, W.A., 1966. Habituation: A model phenomenon for the study of neuronal substrates of behavior. Psychol. Rev., 73, 16–43. 10.1037/h0022681

Urick, R.J., 1975. Principles of Underwater Sound. McGraw-Hill, New York. [Online]. Available: https://books.google.com [Accessed 24 March 2025]

Varello, R., Testa, C., Prospathopoulos, A., Greco, L., Asnicar, D., Boaga, J., Cima, F., 2023. Behavioural responses to ultrasound antifouling systems by adult solitary ascidians. J. Mar. Sci. Eng., 11, 1115. 10.3390/jmse11061115

Wakai, M.K., Nakamura, M.J., Sawai, S., Hotta, K., Oka, K., 2021. Two-round Ca^2^⁺ transient in papillae by mechanical stimulation induces metamorphosis in the ascidian *Ciona intestinalis* type A. Proc. R. Soc. B, 288, 20203207. 10.1098/rspb.2020.3207

Wang, M., Moin, P., 2000. Computation of trailing-edge flow and noise using large-eddy simulation. AIAA J., 38, 2201–2209. 10.2514/2.895

Wei, Y., Duan, Y., An, D., 2022. Monitoring fish using imaging sonar: Capacity, challenges and future perspective. Fish Fish., 23, 1347–1370. 10.1111/faf.12693

White, K.N., Edwards, C., Ambrosio, L.J., 2021. Anthropogenic sound in the sea: Are ascidians affected? Gulf Caribb. Res., 32, 1–7. 10.18785/gcr.3201.02

